## supplementary for "Analysis of Inositol Phosphate Metabolism by Capillary Electrophoresis Electrospray Ionization Mass Spectrometry (CE-ESI-MS)"

c) Leibniz-Forschungsinstitut für Molekulare Pharmakologie, Robert-Rössle-Str. 10, 13125 Berlin, Germany.

d) Institute of Crop Science and Resource Conservation, Department of Plant Nutrition, Rheinische Friedrich-Wilhelms-University Bonn, 53115 Bonn, Germany.

e) Signal Transduction Laboratory, National Institute of Environmental Health Sciences, National Institutes of Health, Research Triangle Park, NC 27709 USA

f) CIBSS - Centre for Integrative Biological Signalling Studies, University of Freiburg, 79104 Freiburg, Germany.

*

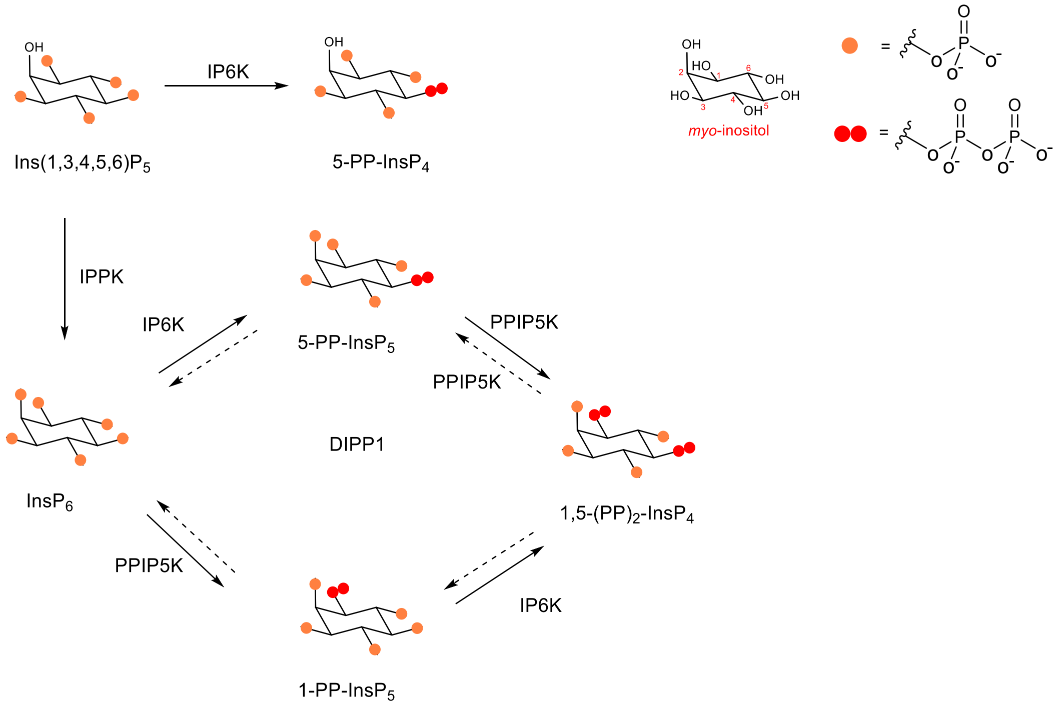


**Supplementary Figure 1** Metabolism of inositol pyrophosphates in mammals. IPPK: inositol pentakisphosphate 2-kinase, IP6K: inositol hexakisphosphate kinase, PPIP5K: diphosphoinositol pentakisphosphate kinase, DIPP1: diphosphoinositol polyphosphate phosphohydrolase 1.

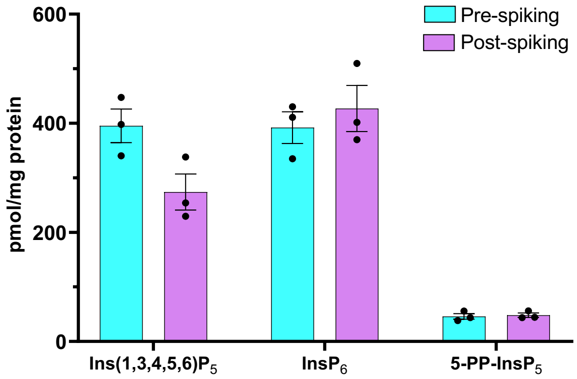


**Supplementary Figure 2** Recovery of 5-PP-InsP and $InsP_6$ from mammalian cell extracts with the $TiO_2$ extraction procedure. 20 µM [$^{13}C_6$]Ins(1,3,4,5,6)$P_5$, 20 µM [$^{13}C_6$]$InsP_6$ and 5 µM [$^{13}C_6$]5-PP-$InsP_5$ were spiked into HCT116$^{NIH}$ cell extracts before the measurement step (post-spiking), or added to culture dishes containing HCT116$^{NIH}$ cells before the perchloric acid extraction step (pre-spiking). The applied InsP purification protocol showed full recovery for 5-PP-$InsP_5$(105%) and $InsP_6$ (109%), and good recovery for Ins(1,3,4,5,6)$P_5$ (69%). Data are means ± SEM from three independent experiments, individual values are shown with dots.

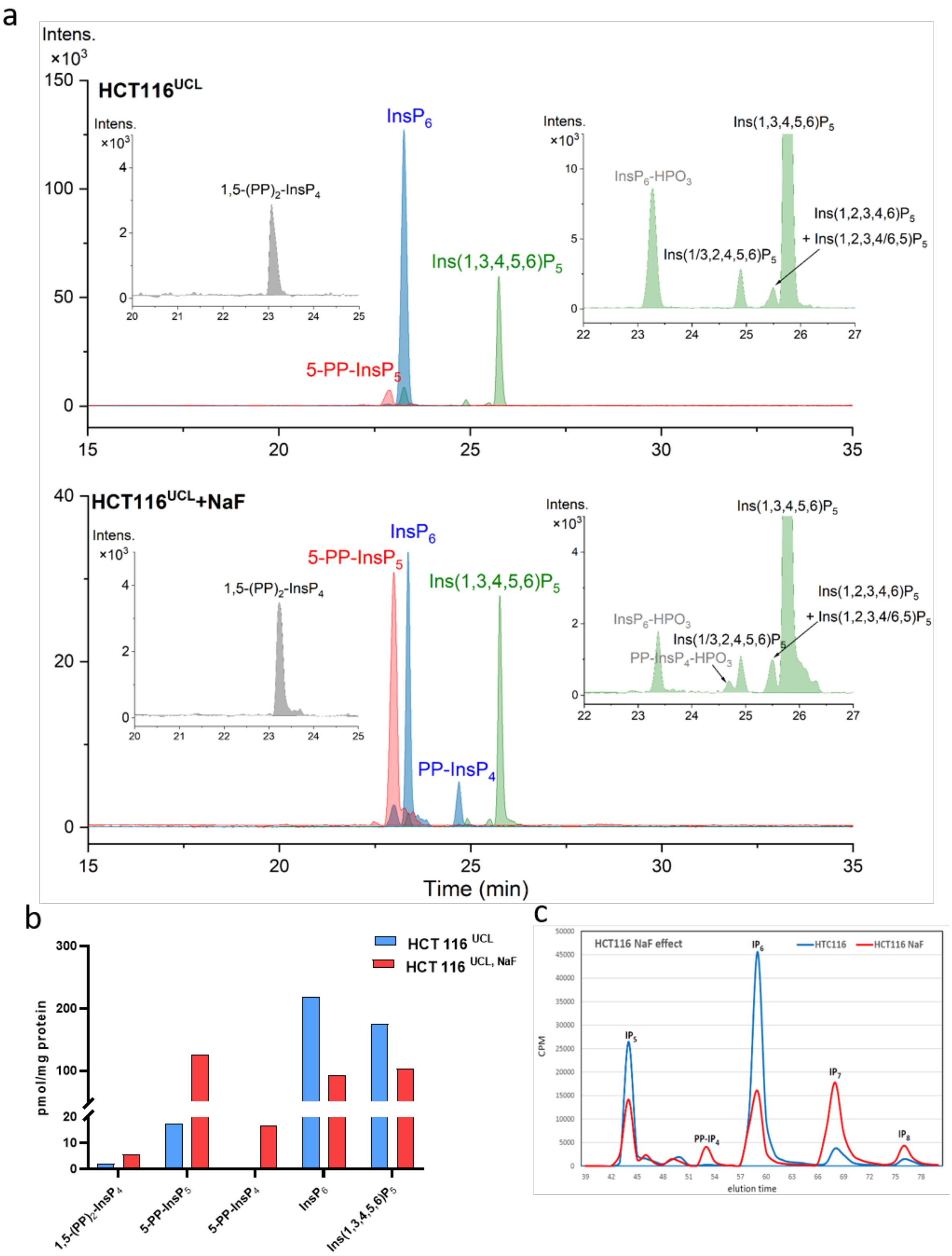


**Supplementary Figure 3** Analysis of inositol (pyro)phosphates in HCT116$^{UCL}$ cells by CE-MS and SAX-HPLC. **(a)** Extracted ion electropherograms (EIEs) of the main inositol (pyro)phosphates in HCT116$^{UCL}$ cells, untreated and after 1 h incubation with 10 mM sodium fluoride (NaF). EIEs of 1,5-(PP)$_2$-InsP$_4$ and minor InsP$_5$ isomers are presented separately. 20

μM [$^{13}C_6$]Ins(1,3,4,5,6)$P_5$, 20 μM [$^{13}C_6$]Ins$P_6$, 10 μM [$^{13}C_6$] 5-PP-Ins$P_5$, 1 μM [$^{13}C_6$] 1-PP-Ins$P_5$ and 1 μM [$^{13}C_6$] 1,5-$(PP)_2$-Ins$P_4$ were spiked. **(b)** CE-ESI-MS results for inositol (pyro)phosphates (amount normalized by protein content) in HCT116$^{UCL}$ cells showing the effect of NaF treatment; **(c)** SAX-HPLC analysis of [$^3$H]-inositol-labeled HCT116$^{UCL}$ cells for NaF treatment.

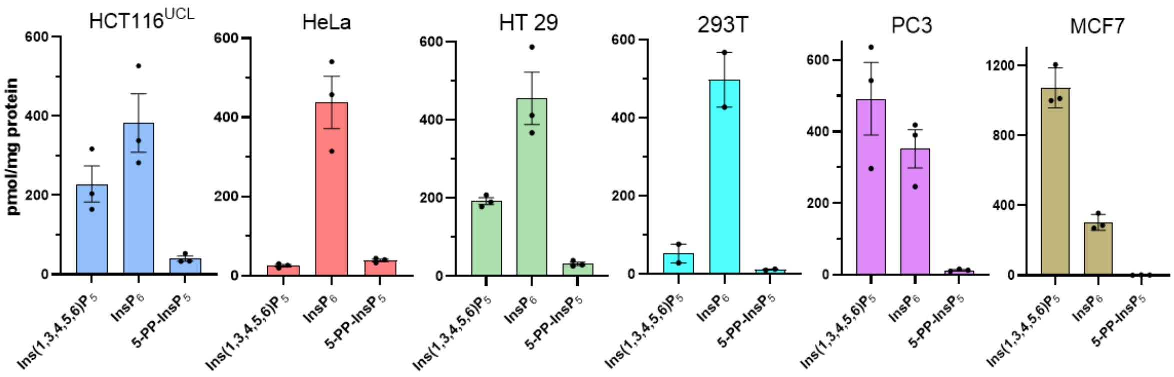


**Supplementary Figure 4** InsP levels normalized by protein content in human cell lines: HeLa, HCT116[UCL], HT29, PC3, 293T and MCF7. The results vary between 25-1080 pmol/mg protein Ins(1,3,4,5,6)$P_5$, 300-500 pmol/mg protein $InsP_6$, and 1-40 pmol/mg protein 5-PP-$InsP_5$ across the cells tested.  Data are means ± SEM from three independent experiments.

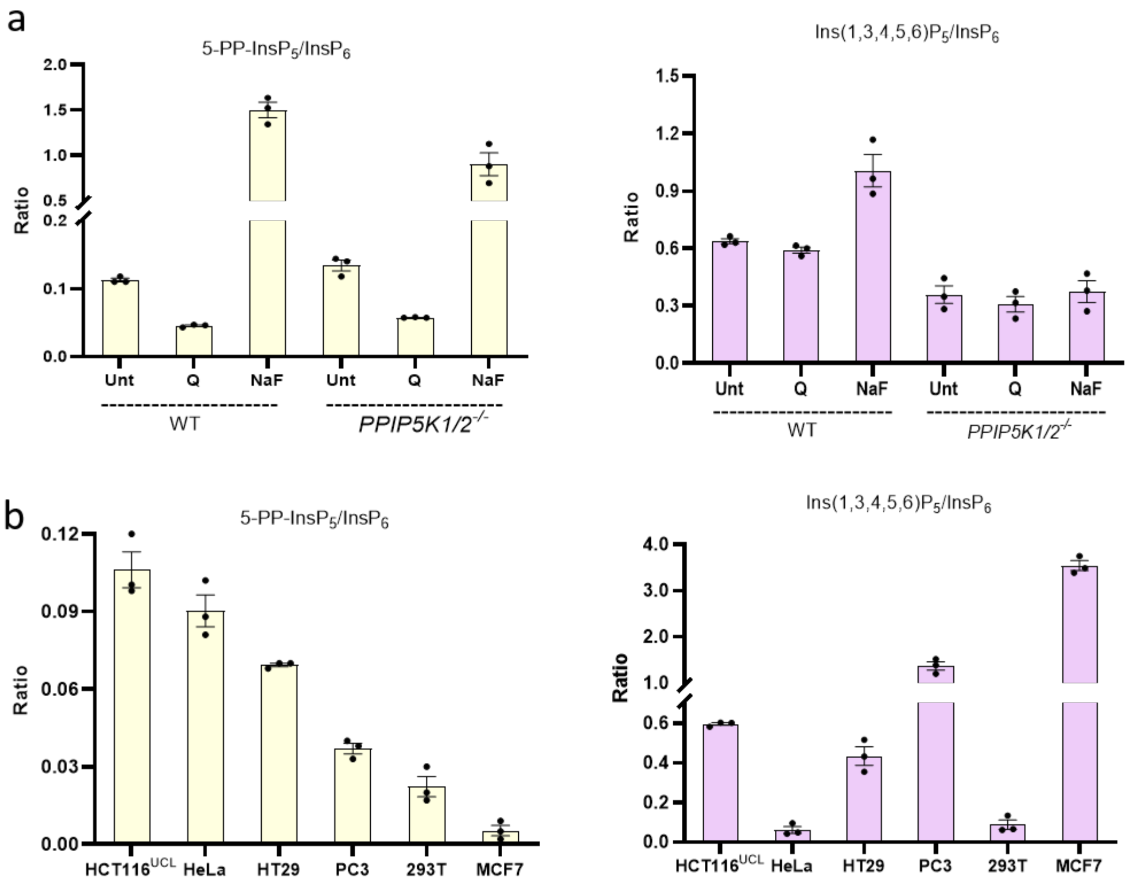


**Supplementary Figure 5** Ratios of InsPs and PP-InsPs in different cell lines. **(a)** Ratio of 5-PP-$InsP_5$/$InsP_6$ (yellow) and Ins(1,3,4,5,6)$P_5$/$InsP_6$ (purple) in HCT116$^{NIH}$ and HCT116$^{NIH}$*PPIP5K*$^{-/-}$ cells after treatment with NaF (10 mM for 60 min) or the inositol polyphosphate kinase (IPMK) inhibitor quercetin (Q; 2.5 μM for 30 min); **(b)** Ratio of 5-PP-$InsP_5$/$InsP_6$ (yellow) and Ins(1,3,4,5,6)$P_5$/$InsP_6$ (purple) in HCT116$^{UCL}$, HeLa, HT29, PC3, 293T and MCF7 cells. Data are means ± SEM from three independent experiments.

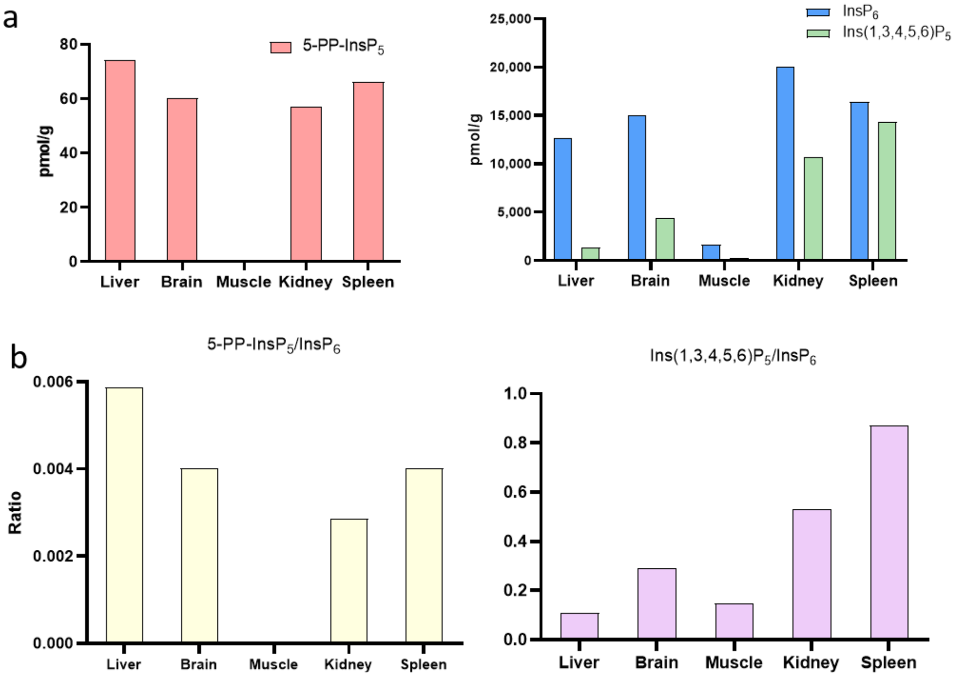


**Supplementary Figure 6** Inositol phosphates and pyrophosphates in organs. **(a)** Amount of inositol (pyro)phosphates in different mouse organs (normalized by organ weight), including liver, brain, muscle, kidney and spleen; **(b)** Ratio of 5-PP-$InsP_5$/$InsP_6$ (yellow) and Ins(1,3,4,5,6)$P_5$/$InsP_6$ (purple) in different mouse organs. Standards of 4 μM [$^{13}C_6$]5-PP-$InsP_5$, 120 μM [$^{13}C_6$]$InsP_6$, and 20 μM [$^{13}C_6$]Ins(1,3,4,5,6)$P_5$ were spiked into organ extracts for quantification.

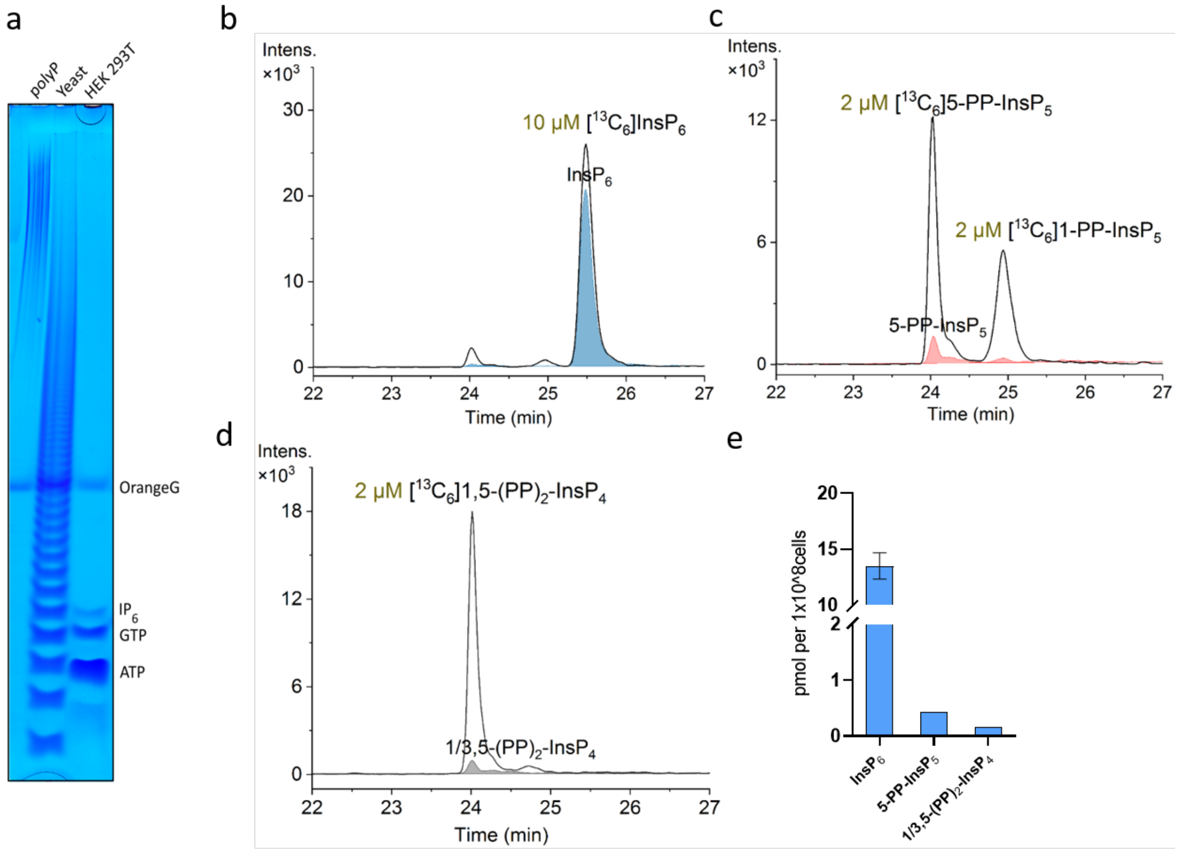


**Supplementary Figure 7** CE-ESI-MS analysis of inositol (pyro)phosphates in wild type *Saccharomyces cerevisiae*. **(a)** Yeast extract analyzed by PAGE. A 33% PAGE gel was loaded with poly standard, $TiO_2$ purified wild type yeast *Saccharomyces cerevisiae* (5 $OD_{600}$) extract and $TiO_2$ purified HEK 293T (10cm plate 80% confluent) extract. After running the phosphate-rich metabolites were visualized by Toluidine staining. The very abundant polyP present in yeast is revealed by the distinctive ladder. The polyP ladder conceals the InsP and nucleotide signals that are conversely easily detectable in the HEK 293T cell extract. The result is representative of an experiment repeated at least four times; **(b-d)** EIEs of $InsP_6$, $InsP_7$ and $InsP_8$ along with spiked 10 µM [$^{13}C_6$]$InsP_6$, 2 µM [$^{13}C_6$]5-PP-$InsP_5$, 2 µM [$^{13}C_6$]1-PP-$InsP_5$, and 2 µM [$^{13}C_6$]1,5-$(PP)_2$-$InsP_4$, respectively; **(e)** Amounts of inositol (pyro)phosphates (normalized by cell counts) in *S. cerevisiae*. Data are means ± SD from independent duplicates

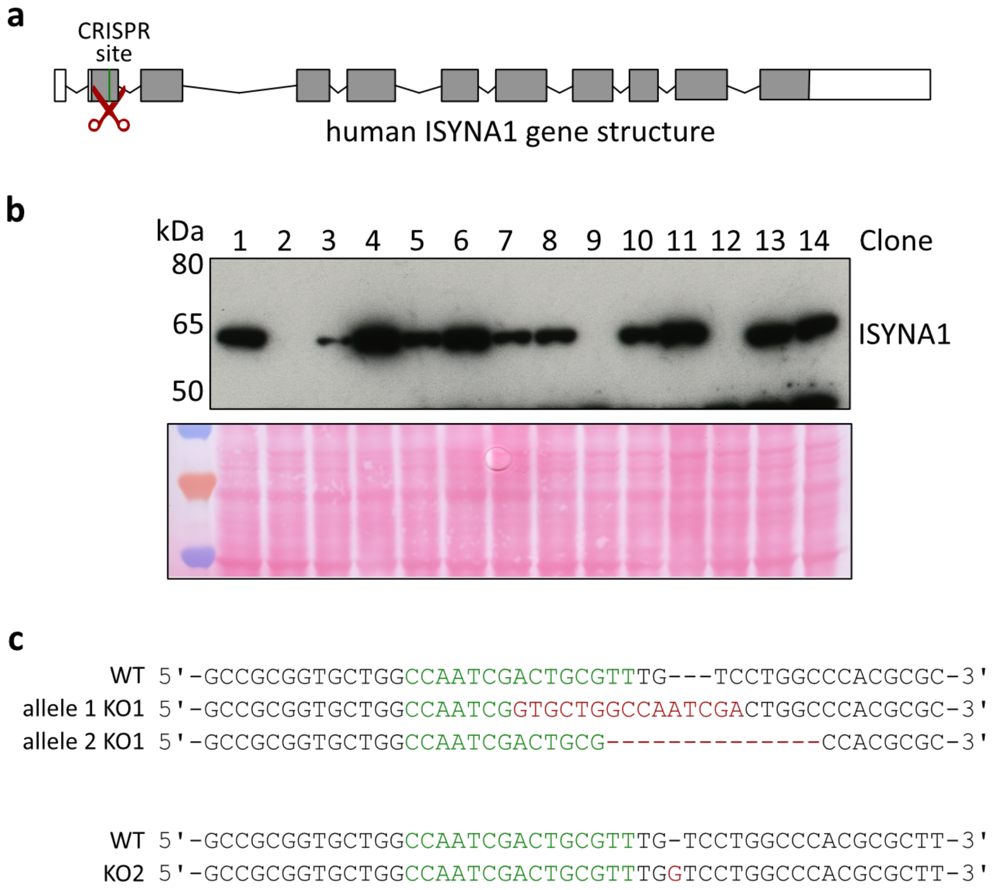


**Supplementary Figure 8** Generation of ISYNA1 knockout clones. **(a)** Schematic structure of the human ISYNA1 gene, with exons shown in grey, indicating the Alt-R CRISPR targeted region; **(b)** Western blotting for ISYNA1 revealed its absence in three of the HCT116$^{UCL}$-derived clones tested. Ponceau red staining confirmed similar protein loading. We chose to focus on two independent clones we called *ISYNA1$^{-/-}$*KO1 and *ISYNA1$^{-/-}$*KO2; **(c)** The mutations around the CRISPR site were determined by PCR and Sanger sequencing. Sequencing found that KO1 has an insertion on one allele and a deletion on the other. All sequences from clone KO2 showed a single nucleotide insertion.

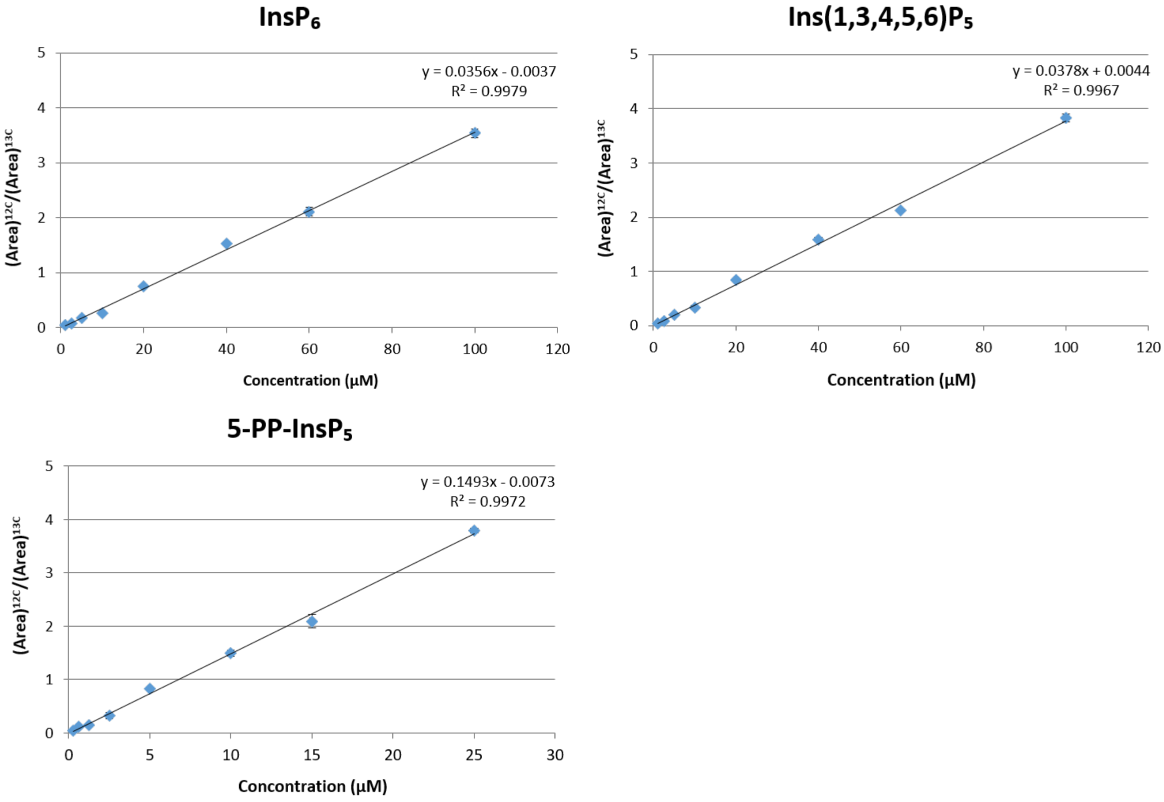


**Supplementary Figure 9** Calibration curves for quantification of Ins(1,3,4,5,6)$P_5$, Ins$P_6$ and 5-PP-Ins$P_5$, in mammalian cells. Standards were spiked with 5 µM [$^{13}C_6$] 5-PP-Ins$P_5$, 20 µM [$^{13}C_6$] Ins$P_6$ and 20 µM [$^{13}C_6$] Ins(1,3,4,5,6)$P_5$. The ratios of peak areas for Ins(1,3,4,5,6)$P_5$, Ins$P_6$ and 5-PP-Ins$P_5$ with respect to internal standards [$(\text{Area})^{12C}/(\text{Area})^{13C}$] are plotted against the concentrations of the standards. Data are means ± SD from duplicates.
